# Using known births to account for delayed marking in population estimation of North Atlantic right whales

**DOI:** 10.1101/2024.10.11.617830

**Authors:** Daniel W. Linden

## Abstract

Population estimation using capture-recapture modeling typically requires that individuals are identifiable by unique marks. North Atlantic right whales (*Eubalaena glacialis*) can be identified by natural callosity patterns on their heads that are established nearly a year after birth, which has facilitated population monitoring using extensive aerial surveys. A well-maintained catalog of individual sightings has been used to annually estimate population size with a Jolly-Seber (J-S) model using a Bayesian state-space framework. Given that young animals cannot enter the catalog before an established callosity pattern, the terminal year population estimate never includes new calves despite breeding area surveys that provide a nearly complete census of births. Here, I illustrate a simple modification to the J-S likelihood whereby the number of expected entrants is a function of known births and a parameter representing initial offspring mortality. A simulation study was used as a proof of concept and indicated increased accuracy and precision of population estimates. The birth-integrated J-S model had more accurate terminal-year estimates of right whale population size that remained consistent during subsequent model fitting to additional years of sightings data. While the bias corrections were fairly small (5%) given low per capita calving rates, the demonstrated improvement in accuracy will be helpful to the conservation and management processes for this endangered species. Integrated modeling approaches make better use of available data and can improve inferences on population dynamics.

## 1 Introduction

Accurate estimates of population size are a fundamental tool for the conservation and management of wild animals (Williams et al. 2002). While many different types of observations and estimation methods can be used for population monitoring, repeated surveys of identified individuals modeled within an open population capture-recapture framework provides one of the best approaches to understand population change over time (Seber and Schofield 2019). The value of individual identification is particularly apparent when a species is otherwise difficult to observe but possesses natural markings, allowing for collection of non-invasive sightings (Hammond et al. 2021). The ability to collect observations of animals without the need for physical capture has greatly improved opportunities for effective and efficient wildlife population monitoring at broad spatial and temporal scales (Stevick et al. 2001).

Traditional capture-recapture models assume that all individuals in a population are equally likely to be captured (or sighted and identified) and that marks used to identify individuals are not lost (Pollock et al. 1990). For many species of mammals, characteristic natural markings are absent or still changing in young animals (Ling 1970), causing these individuals to violate standard model assumptions (Yoshizaki et al. 2009). Advanced modeling techniques can address certain types of mark loss (Conn et al. 2004), variation in mark quality (Stevick et al. 2001, Bonner and Holmberg 2013) and even partial identity (Augustine et al. 2018), typically at the expense of increased uncertainty. A straightforward solution for species with delayed development of a permanent mark is to limit inferences to the mature segment of the population that can be confidently identified. Observations of young animals can be modeled distinctively from identified individuals to provide information on reproduction instead of population size, a strategy often used with integrated population models (Schaub and Kéry 2021).

North Atlantic right whales (*Eubalaena glacialis*; hereafter, right whales) are extensively monitored with aerial and boat surveys throughout their range in the western Atlantic Ocean, providing an unprecedented catalog of photographed individuals since the 1980s (Hamilton et al. 2007). An open population capture-recapture model has served as the primary tool for right whale population assessment on an annual basis in recent years (Pace et al. 2017), though many additional efforts to monitor individual health and document mortality and calving events have been made possible by a consortium of nonprofit science organizations, academic institutions, and government agencies (https://www.narwc.org/). The ability to identify individual right whales is enabled by unique natural callosity patterns on the head, which become permanent within a year after birth (Kraus et al. 1986). A nearly complete census of right whale calving events in the southeastern United States (Keller et al. 2012) provides many observations of young individuals that cannot be uniquely identified until later sightings and/or genetic information (Frasier et al. 2007) allows some linkage to the birth.

While upwards of 60% of individual right whales end up having a known birth year, the delayed marking results in the absence of calves from the terminal year of the annually estimated population size.

Here, I present a specific modification to the Jolly-Seber (J-S) model (Jolly 1965, Seber 1965) to integrate known births into the estimation of population size. The J-S model uses observed capture or sighting histories to estimate probabilities of individual entry (i.e., birth or immigration) and exit (i.e., death or emigration) from a population, while accommodating the potential for imperfect detection (i.e., false negatives) of individuals during surveys. Population size is then determined by the total individuals estimated to be alive in a given year, whether or not they are sighted. The entry probabilities are typically determined by the individual capture histories alone, which presents a lost opportunity when additional information on the expected entries (e.g., observed births) is available. I first briefly describe the multistate formulation of a J-S model in a hierarchical Bayesian framework (Kéry and Schaub 2012) and how the likelihood is easily modified to integrate known births. I then test the new model with a small simulation study as a proof of concept. Finally, I apply the integrated model to sightings of North Atlantic right whales to illustrate the improved estimates of population size that result from incorporating the annual calf counts.

## 2 Methods

### 2.1 Jolly-Seber model

Many versions of the J-S model have been described and Kéry and Schaub (2012) present several popular options and outline similarities and differences between the available parameterizations. Regardless of the formulation, a hierarchical state-space perspective explicitly separates the ecological state processes (i.e., population dynamics over time) from the observation processes. Individuals transition between various states according to ecological parameters while the conditional observation of those states is determined by detection parameters.

Here, we consider that a population of individuals across time can be in 1 of 3 ecological states: (1) not yet entered the population; (2) alive; and (3) dead. The latent state for *i* = 1, …, *n* individuals across *t* = 1, …, *T* years is represented in the state matrix, *z*_*i,t*_. The observed capture history matrix, *y*_*i,t*_ indicates whether an individual was seen (*y*_*i,t*_ = 1) or not seen (*y*_*i,t*_ = 2) in a given year. To accommodate the possibility for individuals that were alive but never seen, data augmentation is used to expand the observed capture histories with some number, *n*_0_, of pseudo-individuals with all-zero capture histories (Royle and Dorazio 2012). The matrices of observed data and latent states now have values for *n* + *n*_0_ = *M* individuals across *T* years.

For the multistate formulation, all individuals start out as “not yet entered” (*z*_*i*,0_ = 1) for a dummy occasion before *t* = 1. Transitions across years are then dictated by a state transition matrix:

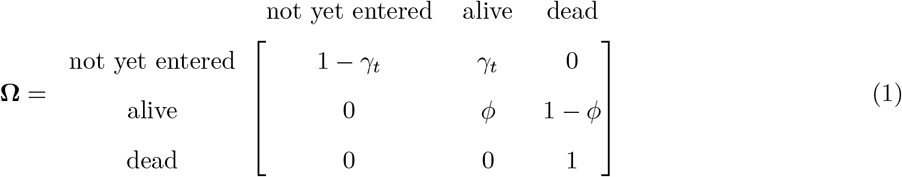

where *γ*_*t*_ is the probability of population entry and *ϕ* is the probability of survival. The latent entries of the *n* observed individuals into the population occur during or before the years they are first observed, while only a small portion of the *n*_0_ pseudo-individuals will be predicted to have ever entered (and been unobserved). Once an individual enters the population, it remains in the alive state according to *ϕ* until it transitions permanently to the dead state. Note that observations of death are not required. The observation matrix specifies the probabilities of observing individuals given their true latent state:

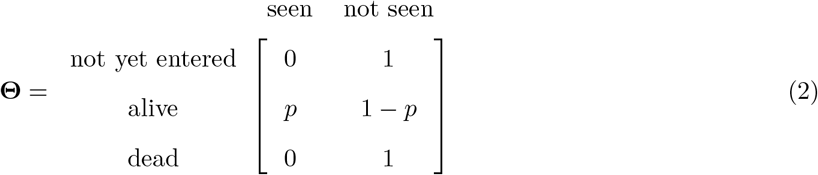

where *p* is the probability of sighting individual *i* in year *t*, given the true state *z*_*i,t*_ = 2.

In this simplest version of the J-S model, the probabilities of survival and sighting are allowed to be constant (no variation across individuals or time) while the probabilities of entry should be year-specific. Technically, *γ*_*t*_ describes a removal process (Kéry and Schaub 2012) that is a function of the number of *M* individuals that have yet to enter the population. The expected number of individuals to enter the population in *t* = 1 is *E*(*B*_1_) = *Mγ*_1_, which includes both individuals that were already in the population prior to *t* = 1 and actual new entries. For subsequent years, expected entries (*B*_*t*_) are calculated as follows:

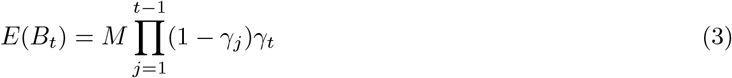

Kéry and Schaub (2012) note that *γ*_*t*_ is a nuisance parameter without ecological interpretation and merely serves to estimate *B*_*t*_. The only data that informs the likelihood in the standard J-S model exists in the capture history matrix, *y*_*i,t*_. Population size (*N*_*t*_) is then estimated by summation of the latent alive states in *z*_*i,t*_.

### 2.2 Partially latent states

There may be information beyond the capture history data that can inform the J-S model about known entries and exits. During model fitting, known latent states (i.e., when values of *z*_*i,t*_ can be indicated with certainty) can be included as data to improve the efficiency of parameter estimation (Kéry and Schaub 2012). Any individuals that contain information about the year of their birth or death could have their corresponding values for *z*_*i,t*_ indicated instead of estimated, reducing uncertainty about probabilities of transition from *z*_*i,t*_ = 1 to *z*_*i,t*_ = 2, and from *z*_*i,t*_ = 2 to *z*_*i,t*_ = 3.

### 2.3 Integrating known births

If the population under study can be assumed to have no immigration, all entries in years *t >* 1 would result from births. A comprehensive survey of reproduction might yield counts of births (*b*_*t*_) that would directly inform the number of expected entries. Assuming the birth counts are collected before the capture surveys, it might be necessary to scale the counts to account for early offspring mortality (*κ*), such that *E*(*B*_*t*_) ≡ *b*_*t*_(1 −*κ*) for all *t >* 1. The following equation can then be used to calculate the new removal entry probability:

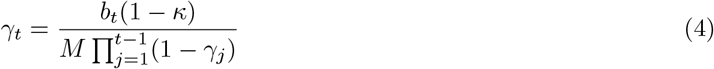

In this manner, the estimate of *γ*_1_ is determined by the capture history data as in the standard J-S model, but all *γ*_*t*_ that follow are a deterministic function of the observed births scaled by offspring mortality.

### 2.4 Simulation study

To assess the accuracy of the birth-integrated J-S model, I simulated random populations and sightings data that approximated most attributes of the right whale survey. The simulated populations all started with *B*_1_ = 250 individuals that transitioned across *T* = 20 years with a constant survival probability of *ϕ* = 0.95 and sighting probability of *p* = 0.8. Births each year followed a Poisson distribution with *λ* = 15. Individuals that entered the population in the terminal year were assigned a sighting probability of *p* = 0.

I designed several scenarios to examine the influence of variation in expected entries and information on known states by varying the amount of offspring mortality (*κ*) and the proportion of individuals with a known age. For the *κ* scenarios, I set 3 different levels: 0.1, 0.3, and 0.5. The variation in *κ* resulted in average expected entries (*B*_*t*_) of 13.5, 10.5, and 7.5 individuals each year, approximating the per-capita birth rates (3–5%) of right whales. For the known-age scenarios, I explored the influence of having known ages for 0% and 60% of all observed individuals in the simulated data. For each combination of *κ* level and known-age proportion, I simulated 100 capture-recapture data sets and augmented the capture histories with *n*_0_ = 200 pseudo-individuals. Observations of individuals that entered the population in the terminal year were switched to “not seen” to mimic the lack of identifiable marks in newly entered right whales (i.e., calves).

For each data set from the 6 scenarios, I fit both the standard J-S model and the birth-integrated J-S model. All parameter estimation was achieved using Markov chain Monte Carlo (MCMC) methods with NIMBLE (de Valpine et al. 2017) in R (R Core Team 2022). Vague priors were used for the probabilities that were estimated (*ϕ, p, κ*, and *γ*_1_). Given the model simplicity and high sighting probability, I found that a single chain of 1,000 iterations after a burn-in of 1,000 provided sufficiently converged parameter estimates. Model performance across the 100 simulated data sets was assessed by calculating the relative bias (RB) and root mean squared error (RMSE) of the mean population size 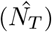 and mean expected entries 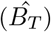,both from the terminal year estimates. I also briefly examined RB of the estimated probabilities, and report 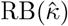,for the birth-integrated model. Finally, I calculated the coefficient of determination (*R*^2^) between actual entries (*B*_*t*_) and estimated entries 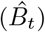,excluding the first year (*t* = 1), when many individuals are already in the population, and the terminal year (*t* = *T*), when new births are known to be unobserved. An *R*^2^ value was calculated for each of the 100 simulated data sets per scenario, and I report the mean and standard deviation.

### 2.5 Case study: Retrospective population sizes of North Atlantic right whales

Pace et al. (2017) describe the J-S model used to estimate population size since 1990 for North Atlantic right whales in the western Atlantic. Individuals are identified primarily by natural markings (Hamilton et al. 2007) with additional information from genetic sampling (Frasier et al. 2007). A photo-identification database and whale catalog curated by the New England Aquarium (NEAq) on behalf of the North Atlantic Right Whale Consortium (NARWC) is updated periodically as evidence is assessed and individual identification is confirmed; the confirmation process can take upwards of a year following collection of a sighting. Sightings are aggregated by individual into survey years (1 December-November 30) to align with the calving season and the seasonal distribution of survey effort, and a population estimate is produced by October each year for sightings from 1990 through the previous year. Calves born in the terminal year are missing from the catalog given their lack of identifying features.

During the J-S model fitting, known alive states are assigned for all survey years between the first and last years with sightings, including for individuals first seen prior to 1990. In addition to sightings of live and dead individuals, those with known birth years are assigned a known state of *z*_*i,t*_ = 1 for all survey years prior to birth. Any years with equivocal evidence of the true state for an individual are assigned as unknown (*z*_*i,t*_ = NA).

Age and sex are known for >60% and >90% of individuals, respectively. Given a known birth year, individuals are classified each year into 1 of 6 age classes (0,1,2,3,4,5+) to accommodate modeling variation in survival for younger animals. Pace et al. (2017) originally assigned the age at entry to be 5+ (adult age class) for individuals with an unknown birth year (mostly whales born before 1990), though other options exist for explicit handling of age (Hostetter et al. 2021). Generalized linear models are used to estimate variation in survival and sighting probabilities, including fixed effects for age and sex. Random effects are used to accommodate unexplained variation across years and individuals for sighting probability, and years for survival probability. Data and code for the latest North Alantic right whale population estimate (Linden 2023) is publicly available (https://github.com/NEFSC/PSD-NARW_popsize).

I examined the performance of the birth-integrated J-S model vs. the standard J-S model by comparing several terminal-year population size estimates (2019, 2020, 2021) as they would have been estimated according to the annual assessment cycle. As a benchmark, I used the latest population estimate from sightings through 2022 (Linden 2023), which would be expected to contain more accurate estimates for the preceding years (2019–2021) than those available from each retrospective terminal year. To replicate each terminal-year population estimate as it would have been estimated at the time, I filtered sightings and whale catalog entries to reflect only the information that would have been available. Given the timing of the photographic assessment process by NEAq, this meant all sightings through the terminal year and known state updates for adult female whales seen in the following calving season. For example, the terminal-year estimate for 2019 would include all sightings and cataloged whales through 30 November 2019, with known states updated for adult individuals seen during the 2020 calving season.

For each of the 3 terminal years, I fit both the standard J-S model and the birth-integrated J-S model. Offspring mortality was estimated as a constant (*κ*), though temporal variation might be warranted with sufficient data. The use of both vague prior distributions and known states as data was executed as previously documented (Pace et al. 2017, Linden 2023). Convergence was sufficient with 10,000 iterations across 2 chains after a burn-in of 5,000. As with the simulations, model performance was assessed by calculating the RB and RMSE of the terminal population estimates 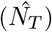, recognizing that the latest estimates (Linden 2023) were a proxy for “truth” and consisted of posterior distributions instead of single values.

## 3 Results

### 3.1 Simulation results

The birth-integrated J-S model improved the accuracy of the terminal-year population estimates compared to the standard J-S model across all examined scenarios (Table 1; Figure 1). As expected, neither model exhibited bias in estimation of *ϕ* and *p* (Table S1), but the standard model consistently underestimated the expected entries in the terminal year (*B*_*T*_ ; Table 1; Figure S1). Errors were greater at lower offspring mortality rates (Table 1), when there were more expected offspring that would be missing from the capture histories. The RMSEs of 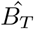 for the standard model matched the expected entries of 13.5, 10.5, and 7.5 individuals, respectively, across the 3 *κ* levels.

**Table 1.**
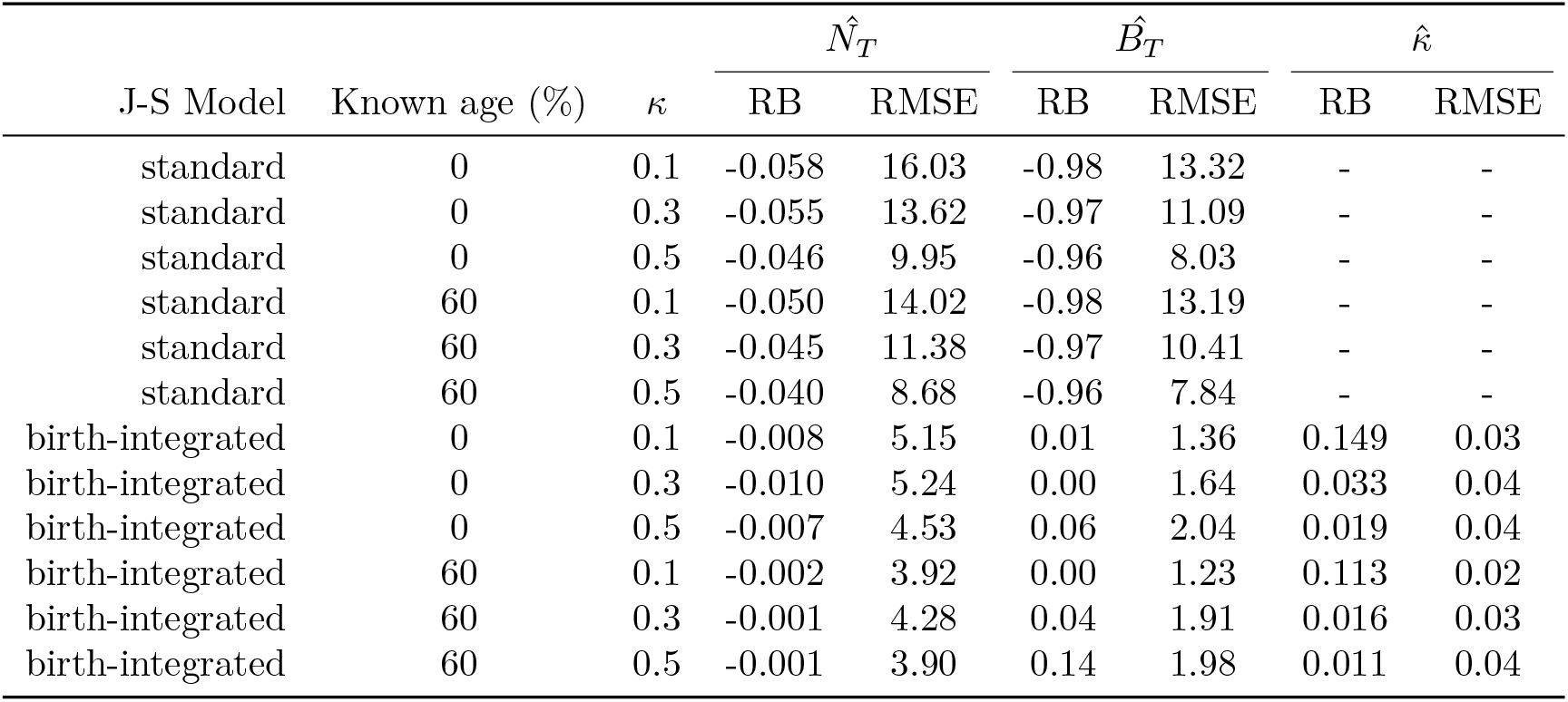
Mean relative bias (RB) and root mean square error (RMSE) for estimates of terminal-year population size 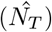 and entries 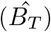 from the standard and birth-integrated Jolly-Seber (J-S) models fit to simulated capture-recapture data. Simulation scenarios varied by percentage of known-age individuals and the specified probability of offspring mortality (*κ*). Additionally, RB and RMSE reported for the estimates of offspring mortality 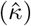 provided by the birth-integrated J-S model.

**Figure 1.**
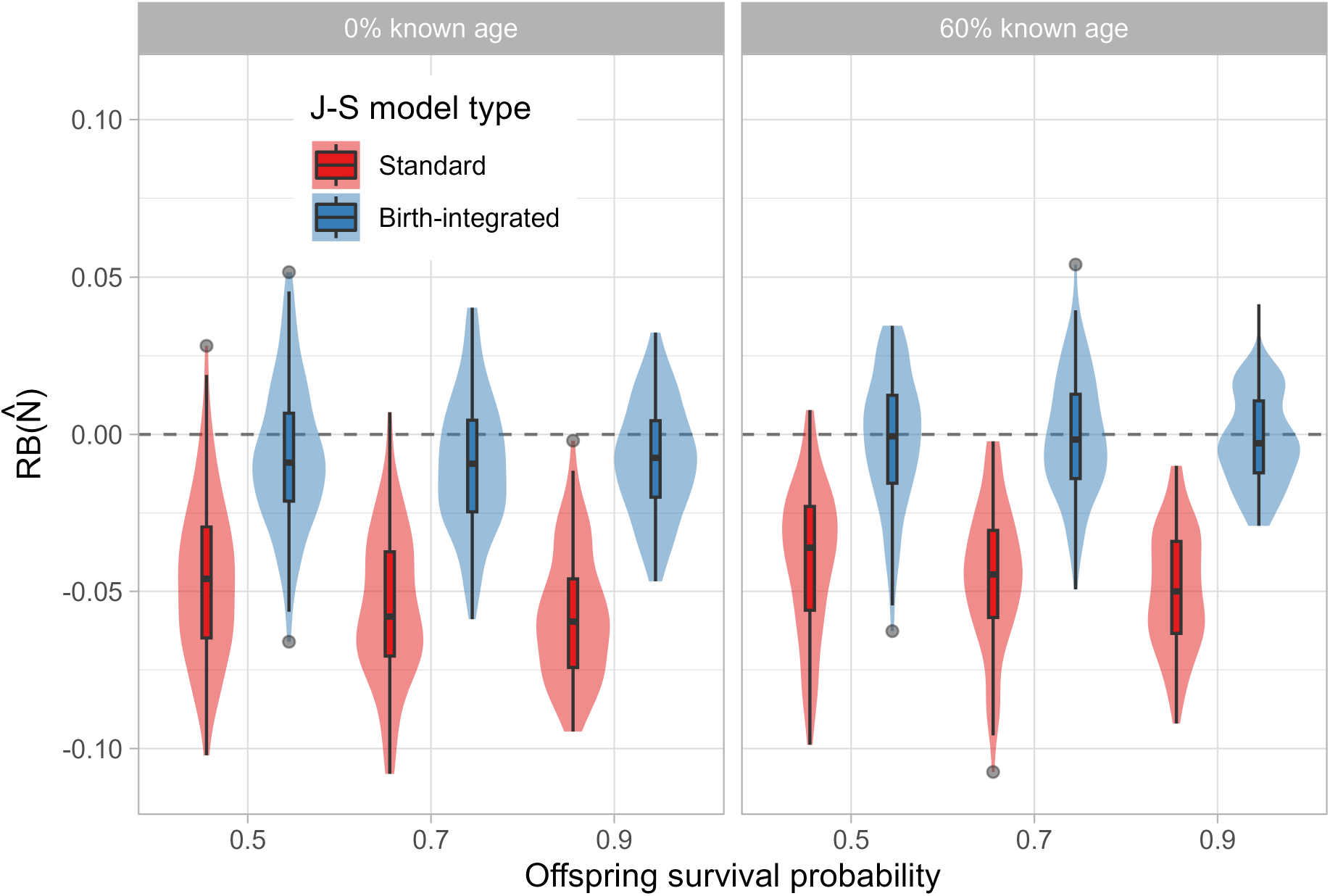
Distributions of relative bias (RB) for estimates of terminal-year population size 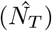 from the standard and birth-integrated Jolly-Seber (J-S) models fit to simulated capture-recapture data. Simulation scenarios varied by percentage of known-age individuals and the specified probability of offspring mortality (*κ*), here represented as survival (1 − *κ*).

Aside from improving the terminal year estimates, the birth-integrated J-S model also improved the predicted entries 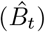 between years 2 and 19 (Table S2). For the standard J-S model with 0% known ages, mean *R*^2^ values ranged from 0.49 to 0.52, while those for the birth-integrated model ranged from 0.68 to 0.79. Increasing the proportion of known ages also improved the correspondence between *B*_*t*_ and 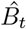. Estimated population sizes 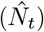 were marginally improved, and only when there were 0% known ages (Table S2). The *R*^2^ values for 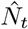 also increased at higher offspring mortality levels (*κ*), likely because these simulated populations exhibited a consistent negative trend on average while populations at *κ* = 0.1 were variably stable.

### 3.2 Improved terminal-year population estimates for North Atlantic right whales

The birth-integrated J-S model improved the correspondence in estimated population sizes between models fit with data filtered through the terminal years 2019–2021 to the most recent estimates available from 2023 (Table 2; Table S3; Figure 2). Differences in 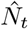 were largely explained by the number of calves observed in the terminal year that were missing from the catalog at the time of estimation (Table S3). Birth-integrated estimates still tended to exhibit a negative bias (Table 2; Figure 2), though current estimates were within the 95% credible intervals. Estimates of calf mortality 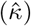 ranged from 0.015 [95% CRI: 0.001, 0.060] for the sightings data through 2019, to 0.035 [0.002, 0.096] for the sightings data through 2021.

**Table 2.**
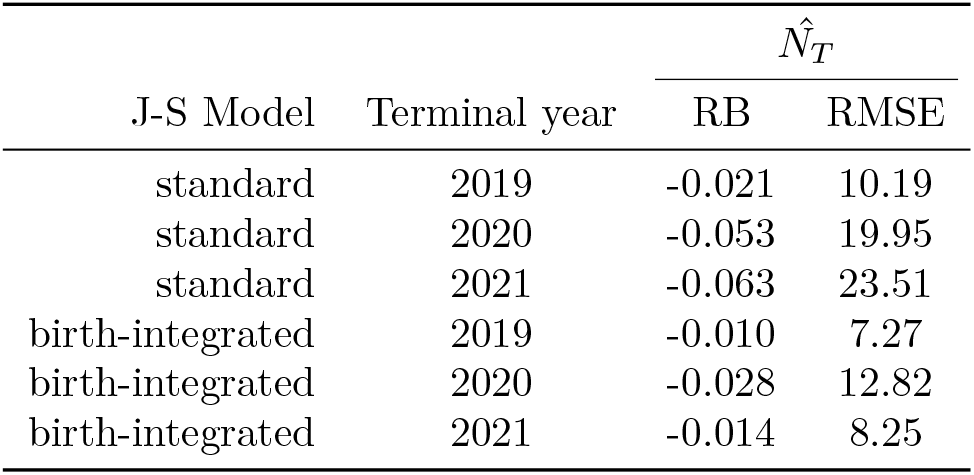
Mean relative bias (RB) and root mean square error (RMSE) for estimates of terminal-year population size 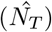 from the standard and birth-integrated Jolly-Seber (J-S) models fit to sightings data of North Atlantic right whales. Data were limited to the information available when each terminal year population size was first estimated. True population sizes were derived from the latest model estimates from 2023.

**Figure 2.**
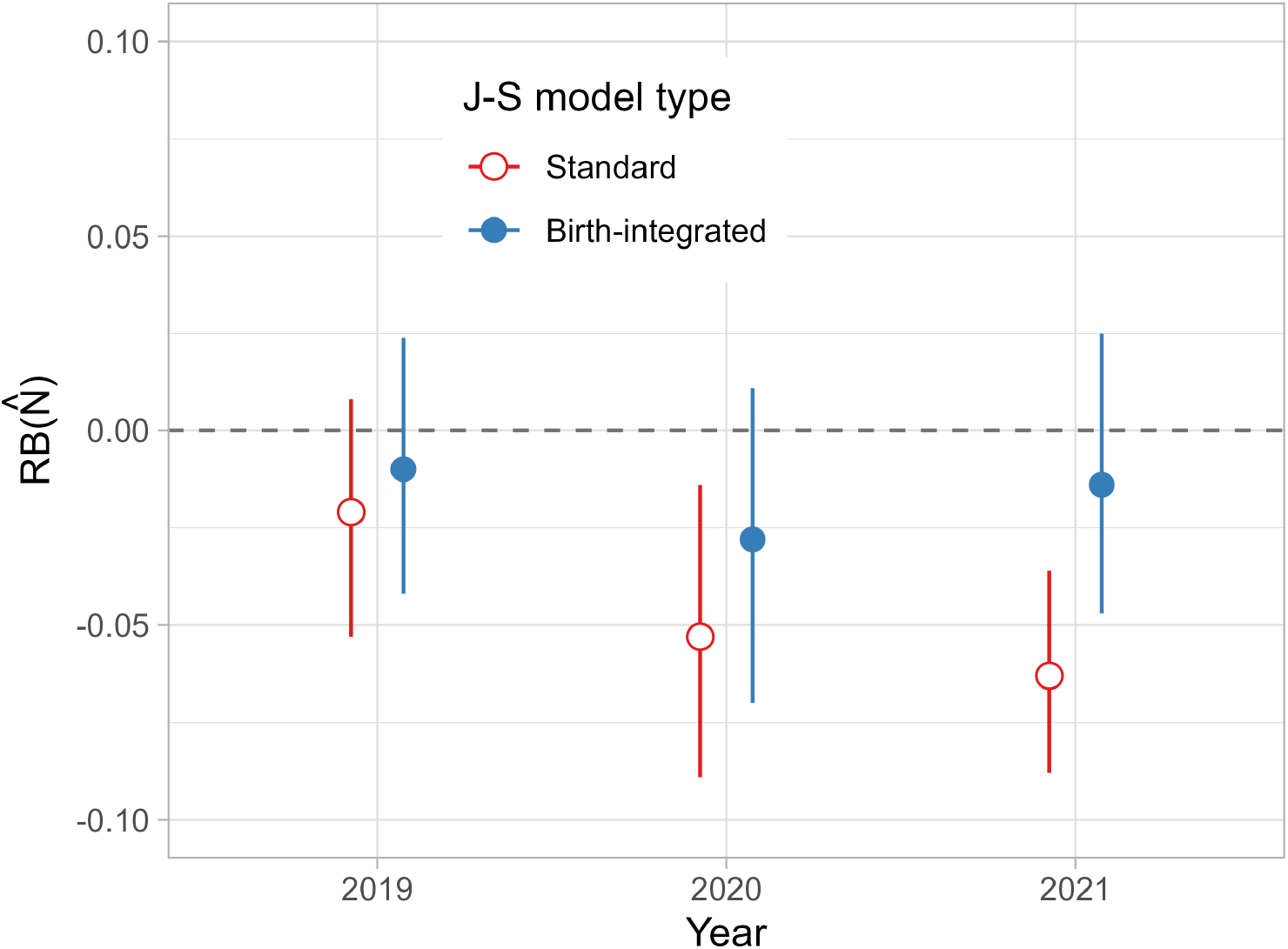
Distributions of relative bias (RB) for estimates of terminal-year population size 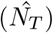 from the standard and birth-integrated Jolly-Seber (J-S) models fit to sightings data of North Atlantic right whales. Data were limited to the information available when each terminal year population size was first estimated. True population sizes were derived from the latest model estimates from 2023.

## 4 Discussion

I developed a birth-integrated version of the J-S model (Jolly 1965, Seber 1965) that allows information on known births to be included in the estimation of population size. This model improves the estimates of annual entries into the population and fixes a specific problem for North Atlantic right whales where delayed marking prevents young whales from being included in the terminal-year population estimate. The modification to the likelihood is straightforward and actually reduces the number of parameters estimated.

North Atlantic right whales are an endangered species that has struggled to persist in a human-dominated landscape (Moore et al. 2021, Runge et al. 2023). Decades of intensive survey effort yielding an unprecedented amount of information on the state of individual whales has indicated that human threats are the primary cause of injury and mortality (Rolland et al. 2016, Knowlton et al. 2022). While annual capture probabilities >0.80 have enabled high precision in right whale population estimates (Pace et al. 2017, Linden 2023), the delayed marking of individuals has resulted in a negative bias in terminal-year population estimates. The magnitude of bias has been a function of the number of calves born in the terminal year. One simple solution might be to add the observed calves in year *t* = *T* to the terminal population estimate 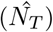, yet early calf mortality could result in such a strategy inducing a positive bias. The birth-integrated J-S model allows for incorporating the known births throughout the time series, increasing model accuracy and correcting the terminal-year estimate.

The comparison of retrospective right whale population estimates with the most current population estimates (Linden 2023) suggested that the birth-integrated J-S model still exhibited a slight negative bias in 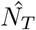. This bias was not apparent in the simulations, indicating that some feature of the right whale sightings data (or model) might be responsible. Despite the callosity pattern developing within a year after birth, there may be further delays in the cataloging of new individuals depending on the available photographs (*P. Hamilton, New England Aquarium, personal communication*). The birth-integrated model induces a sighting probability of 0 for individuals entering the population in the terminal year, but sighting probabilities may also be lower for whales that are 2 or 3 years old. The original Pace et al. (2017) J-S model structure incorporates individual heterogeneity in capture, which can otherwise cause substantial bias in J-S population estimates (Lebreton et al. 1992, Kéry and Schaub 2012). Pace et al. (2017) used simulations to illustrate that the right whale J-S model as constructed was robust to several assumptions regarding capture heterogeneity and unknown age. It should be noted that while the 2023 population estimates are expected to be mostly stable for the retrospective years (2019–2021), the assumption that they represent “truth” for comparison with the birth-integrated J-S model may not hold.

For as much as the birth-integrated J-S model represents a potential improvement to right whale population estimation, additional model complexity might warrant attention. Many other J-S model formulations have been developed (Kéry and Schaub 2012), including versions that explicitly handle known ages and birth information. Hostetter et al. (2021) illustrate an age-structured J-S model that affords improved inferences, particularly with regards to nonlinear relationships between age and mortality risk. Halverson-Duncan (2021) presents a multi-state J-S model that integrates adult and offspring data, though all individuals were physically tagged at time of capture. The Pace et al. (2017) J-S model includes age-based variation in survival probability, though different model structures may be warranted to better accommodate nuance in right whale ecology, including the handling of unknown age individuals. High rates of human-caused mortality likely preclude the potential for inferences on actuarial senescence in North Atlantic right whales, reducing the benefits that might otherwise result from an age-structured version (Hostetter et al. 2021). Other capture-recapture modeling has shown differences in right whale mortality risk by reproductive class and other individual attributes (Reed et al. 2022, Linden et al. 2023), and accommodating this variation could further improve predictions of the true state for some individuals, particularly at the end of the time series when uncertainty regarding unseen whales is highest. For the birth-integrated J-S model, allowing for temporal variation in offspring mortality (*κ*_*t*_) may be worth exploring, particularly given the range in estimates for the retrospective analyses. While such variation would not improve the terminal year estimate, it could provide useful insights regarding threats to right whale population growth and recovery.

Integrated models have become a popular approach for wildlife monitoring when multiple sources of information about a population or species are available (Schaub and Kéry 2021). Combining surveys of reproductive activity with observations of population size (e.g., counts) typically involves specifying a latent population dynamics model that is informed by the time series of available data. The use of such approaches requires cognizance of how inferences are affected when the data being combined are not independent (Abadi et al. 2010, Riecke et al. 2019). Here, the simplicity of a single population without immigration allowed for a small modification to a standard capture-recapture model that improved the accuracy and precision of estimated population processes. Other applications would require similar simplifying assumptions, or additional knowledge of immigration rates that could be incorporated into the estimation of expected population entries. Bayesian hierarchical approaches increase the ease with which new integrated models can be developed, but simulation studies are necessary to increase confidence in the inferences that result.

## Acknowledgements

I am grateful to the NARWC and NEAq for their work in updating and maintaining the Catalog, and for access to the sightings data. The capacity to develop precise estimates of North Atlantic right whale demographic parameters is due to the thousands of photographic captures of whales contributed by hundreds of collaborators working through the NARWC for nearly 40 years. Special thanks to Philip Hamilton for conversations regarding the catalog, and for coordinating data availability. Members of the Atlantic Scientific Review Group graciously reviewed and provided feedback on an earlier version of this manuscript.

## Appendix

**Table S1:**
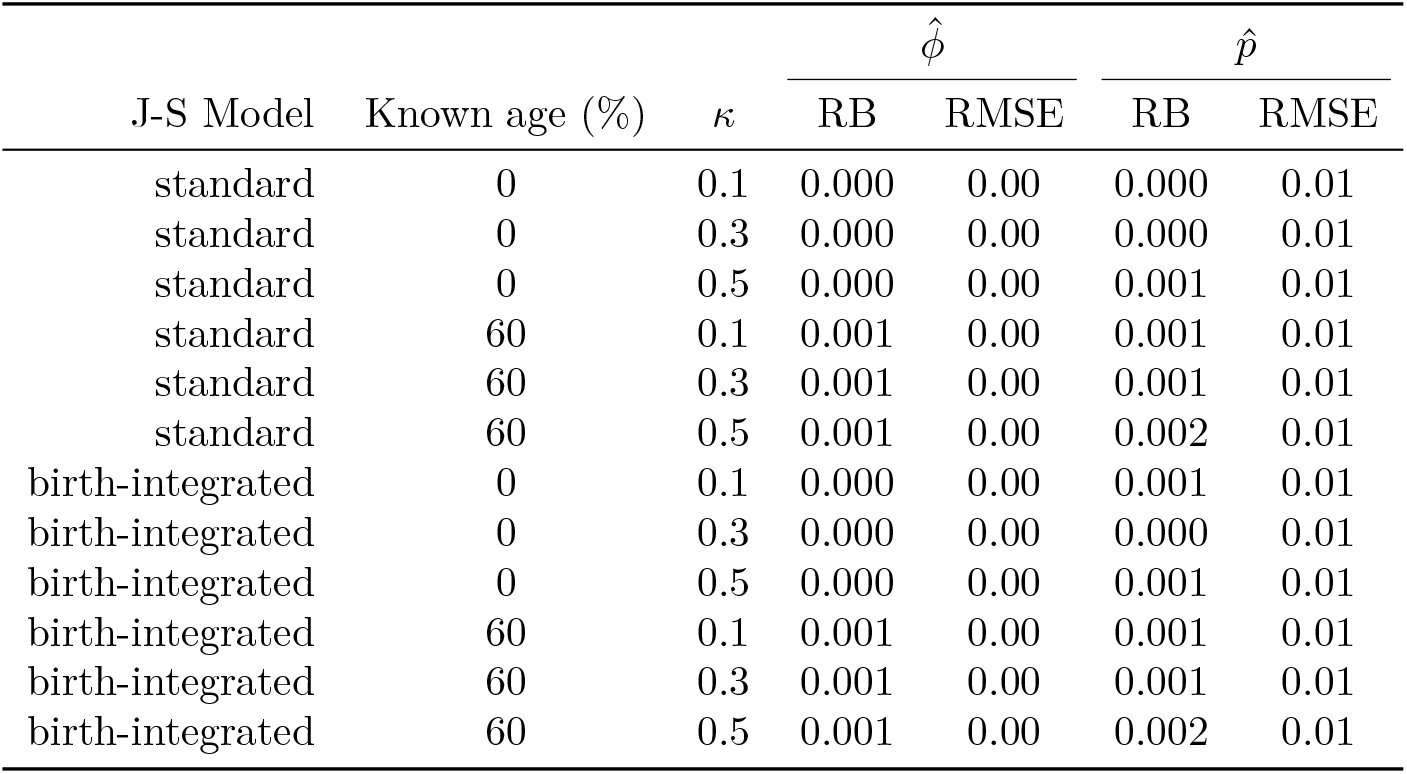
Mean relative bias (RB) and root mean square error (RMSE) for estimates of survival (*ϕ*) and capture probability (*p*) from the standard and birth-integrated Jolly-Seber (J-S) models fit to simulated capture-recapture data. Simulation scenarios varied by percentage of known-age individuals and the specified probability of offspring mortality (*κ*).

**Figure S1:**
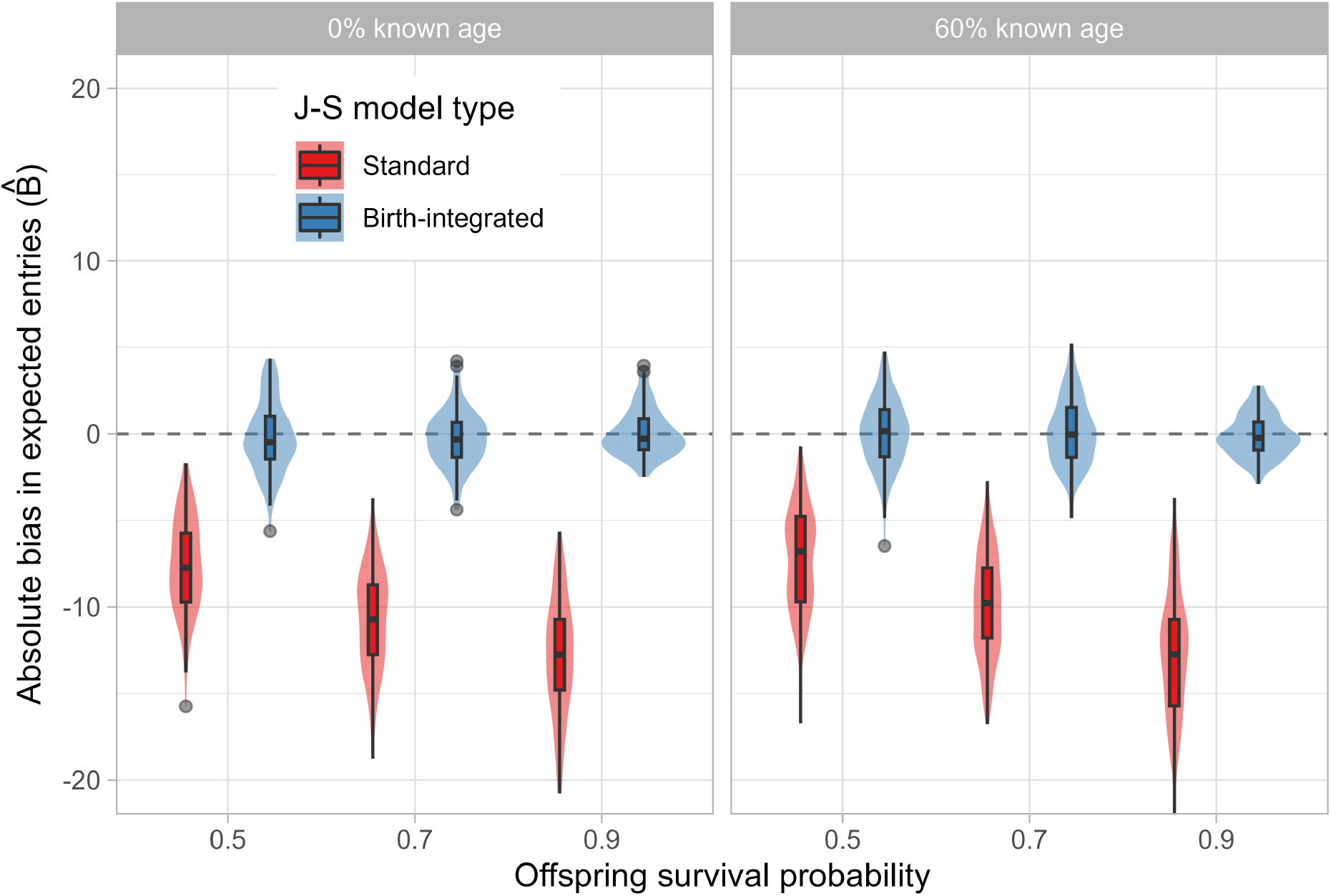
Distributions of absolute bias for estimates of terminal-year entries 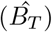 from the standard and birth-integrated Jolly-Seber (J-S) models fit to simulated capture-recapture data. Simulation scenarios varied by percentage of known-age individuals and the specified probability of offspring mortality (*κ*), here represented as survival (1 − *κ*).

**Table S2:** Mean and standard deviation of the coefficients of determination (*R*^2^) between actual entries (*B*_*t*_) and estimated entries 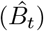 from the standard and birth-integrated Jolly-Seber (J-S) models fit to simulated capture-recapture data. Simulation scenarios varied by percentage of known-age individuals and the specified probability of offspring mortality (*κ*). Comparions excluded the first year (*t* = 1) and terminal year (*t* = 20).

| J-S Model | Known age (%) | $\kappa$ | $\hat{N}_t$ | | $\hat{B}_t$ | |
| --- | --- | --- | --- | --- | --- | --- |
| | | | $R^2$ mean | $R^2$ SD | $R^2$ mean | $R^2$ SD |
| standard | 0 | 0.1 | 0.82 | 0.14 | 0.52 | 0.19 |
| standard | 0 | 0.3 | 0.90 | 0.09 | 0.49 | 0.21 |
| standard | 0 | 0.5 | 0.98 | 0.01 | 0.49 | 0.22 |
| standard | 60 | 0.1 | 0.92 | 0.07 | 0.81 | 0.10 |
| standard | 60 | 0.3 | 0.93 | 0.07 | 0.81 | 0.11 |
| standard | 60 | 0.5 | 0.99 | 0.01 | 0.79 | 0.11 |
| birth-integrated | 0 | 0.1 | 0.87 | 0.11 | 0.79 | 0.11 |
| birth-integrated | 0 | 0.3 | 0.93 | 0.07 | 0.75 | 0.12 |
| birth-integrated | 0 | 0.5 | 0.99 | 0.01 | 0.68 | 0.15 |
| birth-integrated | 60 | 0.1 | 0.93 | 0.06 | 0.86 | 0.07 |
| birth-integrated | 60 | 0.3 | 0.94 | 0.07 | 0.86 | 0.09 |
| birth-integrated | 60 | 0.5 | 0.99 | 0.01 | 0.83 | 0.09 |

**Table S3:** Posterior distributions of estimated North Atlantic right whale population size using both standard and birth-integrated Jolly-Seber (J-S) models. Old model estimates using sightings data that were available at the time of estimation, while the current model estimates are from 2023.

| Year | Calves | Standard J-S (old) |  | Birth-integrated J-S (old) |  | Standard J-S (current) |  |
| --- | --- | --- | --- | --- | --- | --- | --- |
| | | $N_t$ median | 95% CI | $N_t$ median | 95% CI | $N_t$ median | 95% CI |
| 2019 | 7 | 370 | (359,381) | 375 | (362,387) | 378 | (376,382) |
| 2020 | 10 | 337 | (325,350) | 345 | (331,360) | 356 | (352,360) |
| 2021 | 18 | 341 | (333,350) | 358 | (347,370) | 364 | (360,369) |

## Code

R code for the birth-integrated Jolly-Seber model, compatible with BUGS/JAGS/NIMBLE.

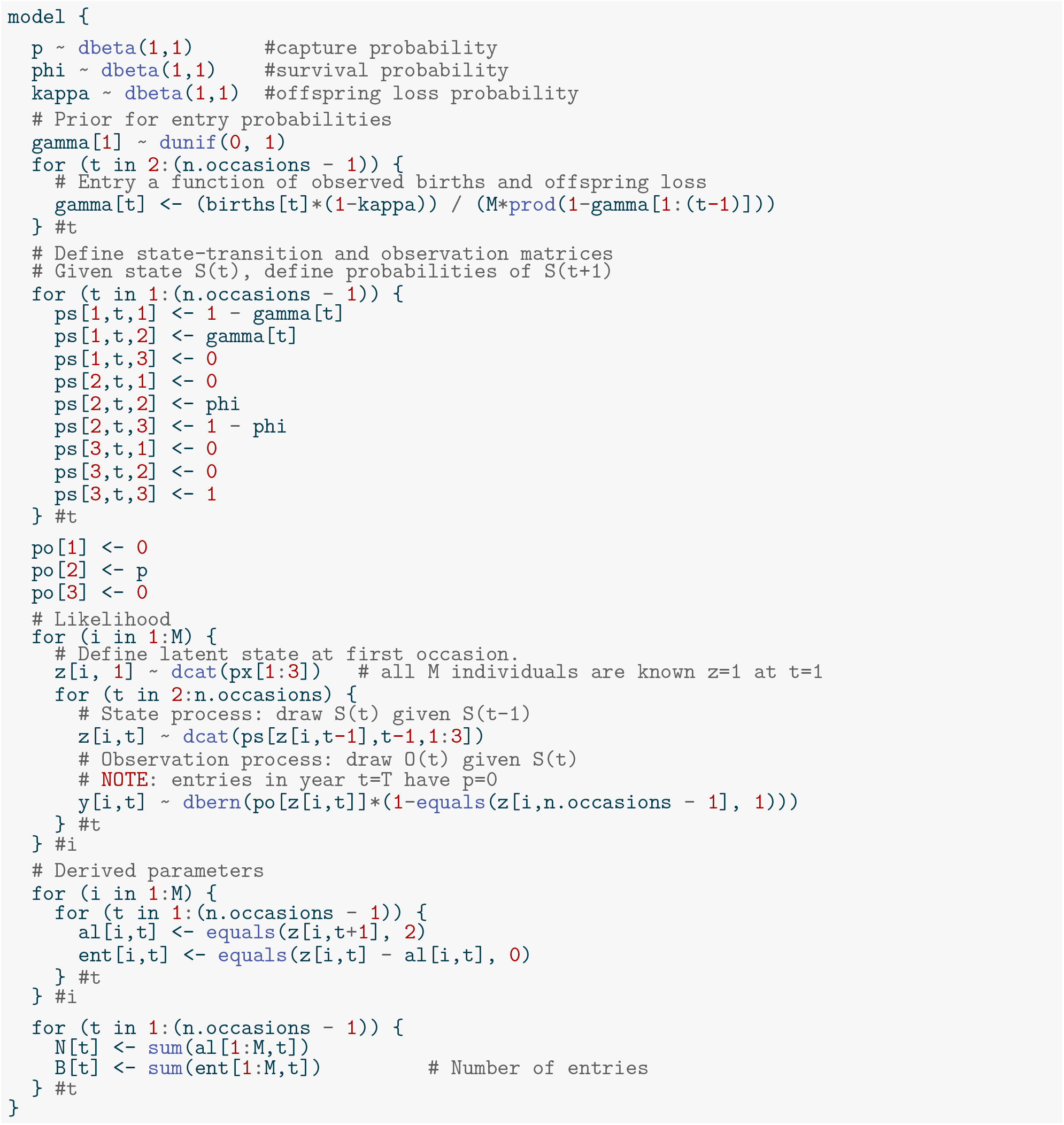

## Notes

### Competing Interest Statement

The authors have declared no competing interest.

